# Enhancing affinity of neutralizing SARS-CoV-2 nanobody through facile structure-guided mutations in CDRs

**DOI:** 10.1101/2024.05.13.593833

**Authors:** Vishakha Singh, Mandar Bhutkar, Shweta Choudhary, Sanketkumar Nehul, Rajesh Kumar, Jitin Singla, Pravindra Kumar, Shailly Tomar

**Author notes:** Corresponding author address: **Shailly Tomar**, Professor, Department of Biosciences and Bioengineering, Indian Institute of Technology Roorkee, Uttarakhand, India 247667.

## Abstract

The optimization of antibodies to attain the desired levels of affinity and specificity holds great promise for development of the next generation therapeutics. This study delves into the refinement and engineering of CDRs through *in silico* affinity maturation followed by binding validation using ITC and pseudovirus-based neutralization assays. Specifically, it focuses on engineering CDRs targeting the epitopes of RBD of the spike protein of SARS-CoV-2. A structure-guided virtual library of 112 single mutations in CDRs was generated and screened against RBD to select the potential affinity-enhancing mutations. Subsequent biophysical studies using ITC provided insights into binding affinity and key thermodynamic parameters. Consistent with *in silico* findings, seven single mutations resulted in enhanced affinity. The mutants were further tested for neutralization activity against SARS-CoV-2 pseudovirus. L106T, L106Q, S107R, and S107Q generated mutants were more effective in virus-neutralizing with IC_50_ values of ∼0.03 µM, ∼0.13 µM, ∼0.14 µM, and ∼0.14 µM, respectively as compared to the native nanobody (IC_50_ ∼0.77 µM). Thus, in this study, the developed computational pipeline guided by structure-aided interface profiles and thermodynamic analysis holds promise for the streamlined development of antibody-based therapeutic interventions against emerging variants of SARS-CoV-2 and other infectious pathogens.

## Introduction

The development of advanced biologic therapeutics, including monoclonal antibodies (mAbs), single-domain antibodies (sdAb) or nanobodies (Nb), and engineered proteins, revolves around enhancing their ability to effectively bind to the specific molecular targets associated with diseases, commonly referred to as antigens. The term “Bio-better” is defined for the biotherapeutic molecules, such as antibodies or proteins, that have been improved to exhibit superior characteristics compared to their original or conventional counterparts ^1^. These improvements could be in terms of efficacy, safety, tolerability, or other desirable properties, presenting a promising strategy for the development of next generation therapeutics ^1,2^. The impeccable specificity and high affinity of antibodies are increasingly exploited for therapeutic and diagnostics purposes ^3–5^. In the process of discovery and development of antibodies, immunization of animals and various display methods (phage, yeast, and ribosome) serve as fundamental techniques ^6,7^. However, reaching the desired levels of affinity and specificity against targeted antigens using traditional approaches may not be consistently successful. Thereby, affinity maturation becomes a valuable strategy to further optimize the binding affinity of antibodies. In nature, this process occurs naturally in the B-cells, in which the host immune system generates high-affinity antibodies against specific antigens through somatic hypermutation ^8,9^. Several *in vitro* techniques based on directed evolution aim to mimic this natural process. However, this daunting task of enhancing the affinity using *in vitro* methods is rather complicated, expensive, and time-consuming. Structure-based computational engineering offers an attractive alternative strategy for the development of biotherapeutic antibodies ^10–12^.

Affinity maturation can be expedited and simplified through computational methods, specifically by employing random virtual mutagenesis guided by the structure-aided interface profiles of antibodies and antigens. This multifaceted task includes structure determination/prediction, identification of binding interface, and calculation of mutational energy changes. The availability of high-quality structural data of antigen and antibody complexes serves as a crucial reference point for the *in silico* affinity maturation process ^13^. The antigen-binding specificity of antibodies is primarily dictated by paratopes, the specialized regions on the molecule responsible for interaction with antigens. Within these paratopes, CDRs (Complementarity-determining regions) play a pivotal role in diversifying and optimizing antibody affinity. Conversely, the remaining framework regions (FRs) contribute significantly to the overall structural integrity of the antibody molecule ^14–16^. To attain a high binding affinity against antigens, antibodies require surface complementarity in terms of shape, size, and chemical nature to ensure effective recognition and interaction with the target antigens. Shape complementarity relies on the aromatic residues that bring the two surfaces together primarily through Van der Waals and hydrophobic interactions. Specificity and strength, on the other hand, are contributed by the electrostatic interactions i.e., salt bridges formed between charged side chains, along with hydrogen bonds connecting oxygen and/or nitrogen atoms ^17,18^. *In silico* affinity maturation has been successfully applied to develop high-affinity antibody fragments such as fragment antigen-binding region (Fabs), Single-chain variable fragments (scFvs), and Variable Heavy domain of Heavy chain (VHHs) against numerous antigens ^19–24^.

VHHs are the antigen binding domain of only heavy chain antibodies that are derived from the camelid’s animals and are also referred to as nanobodies ^25^. Nanobodies, with their compact dimensions of approximately 15 kDa, offer distinct advantages such as specificity, thermodynamic stability, enhanced target engagement, and deeper tissue penetration compared to traditional antibodies. Notably, their smaller molecular size and increased stability simplify the processes of cloning, expression, and purification, allowing for high yields in bacterial, yeast, and mammalian expression systems ^26–32^. The COVID-19 pandemic has accelerated extensive efforts in the field of development and engineering of neutralizing nanobodies against severe acute respiratory syndrome coronavirus 2 (SARS-CoV-2). Despite extensive vaccine administration drives, concerns persist regarding increased transmissibility, immunological resistance, and waning immunity, potentially leading to more severe infection waves ^33^. A critical initial step of infection involves cell entry of viral particles facilitated by the interaction between the spike protein (S-protein) of SARS-CoV-2 and the angiotensin-converting enzyme 2 (ACE2) receptor on the host cell. The receptor-binding domain (RBD) of S-protein, comprises critical residues binding to ACE2, and emerges as a key player in virus entry, offering an attractive target for suppressing and inhibiting virus infection ^34^. Of the other structural proteins of SARS-CoV-2, the S-protein has been recognized as a prime target for the development of neutralizing antibodies ^35–37^. Numerous studies on therapeutics against SARS-CoV-2 have accentuated the efficacy of nanobodies targeting the RBD of the S-protein ^38–48^. These nanobodies neutralize the virus by either directly competing with ACE2 for the same epitope (Ty1, H11-H4, C5, and H3) or by interacting with the non-RBD region of the SARS-CoV-2 S-protein (F2, C1, and VHH72) ^41–44,49^. The incessant emergence of highly contagious escape variants has reinforced the global threat of SARS-CoV-2, necessitating efficient preventive or therapeutic strategies to mitigate any future risks ^50,51^. Global surveillance efforts utilizing genome sequencing unveiled over 5000 amino acid alterations within the genome of SARS-CoV-2, including substitutions, deletions, and insertions, predominantly concentrated within the S-protein ^52^. This demands the need for quick strategies to upgrade the available vaccines and therapeutics for continued efficacy against emerging variants.

Utilizing insights obtained from the crystal structure of the SARS-CoV-2 RBD in complex with nanobody (H11-H4) (PDB: 6ZBP), computational predictions, and *in vitro* validation were employed to engineer and generate high-affinity mutants of the H11-H4 nanobody (referred to native nanobody in this paper). A comprehensive *in silico*, biophysical characterization, and efficacy testing investigations were conducted on a set of 9 mutants that were screened from an *in house* library of 112 single mutants of native nanobody. Using, Isothermal Titration Calorimetry (ITC), the binding affinities of both native and mutant nanobodies were assessed against RBD, aiming to identify the high affinity binders. Pseudovirus neutralization assay successfully identified four mutants of nanobody with significantly improved neutralizing activity compared to their native counterparts. This integrated approach shows potential in expediting the development of antibody based therapeutic interventions targeting emerging variants of SARS-CoV-2 and other pathogens.

## Material and methods

### Software and hardware used for *in silico* study

Protein structures were retrieved from the RCSB-PDB ^53^, and their IDs are indicated as they appear in the text. For structural visualization, PyMol and Crystallographic Object-Oriented Toolkit (Coot) ^54^ were utilized, and Coot was also used for introducing the mutations in the nanobody. Structural comparison and interaction studies were accomplished using PyMol ^55^ and LigPlot+ ^56^. Docking was performed by protein-protein docking software, HADDOCK 2.4 ^57^. Molecular simulation studies were carried out using Gromacs 2022.2 suite ^58^ on an Ubuntu-based LINUX workstation.

### Comprehensive *in silico* mutagenesis

For the computational studies, the coordinates of SARS-CoV-2 RBD and native nanobody (H11-H4) were extracted from the PDB ID: 6ZBP ^43^. A docked complex of the native nanobody and the RBD was generated using HADDOCK 2.4 ^59^. The complex was used as the reference for all the computational studies and for mapping the interacting residue present between the RBD and native nanobody using PyMol and LigPlot+ software. The coordinates of the native nanobody were further used for *in silico* affinity maturation. The first step of *in silico* affinity maturation involves virtual mutagenesis where single mutations were introduced in the selected residues of the CDRs of the nanobody using Coot software, generating a library of nanobody mutants ^60,61^. In Coot, mutations are introduced individually, allowing us to predict their stereochemical effects. The initial position for the new rotamer is chosen based on the most probable orientation ^54^. The substitution mutation was performed majorly to charged, polar, and aromatic residues. Also, mutations to proline, glycine, and cysteine were avoided owing to the peculiar nature of these amino acids. Subsequently, the 3D structures of these mutants were energy-minimized using the CHARMM 27 force field ^62^ in the Gromacs 2022.2 suite. The generated library of nanobody mutants was screened against RBD (Coordinate extracted from 6ZBP) using HADDOCK 2.4 with default parameters. These docked complexes were further screened, employing criteria such as docking scores and favourable interactions compared to the native nanobody complex.

### Protein-protein docking and MD simulations

High affinity complexes selected from the aforementioned pipeline were subjected to MD simulation studies employing the CHARMM 27 force field using the Gromacs 2022.2 suite ^59^. The complex was solvated with a simple point charge (SPC) water model in the cubic box. To neutralize the protein in the water system, counter ions were added, and further energy minimization was executed through the steepest descent algorithm over 50,000 iteration steps. A two-step equilibration process was executed to the energy minimized system for 100 ps, (i) NVT (constant number of particles, volume, and temperature) equilibration, at a reference temperature of 300 K, and (ii) NPT (constant number of particles, pressure, and temperature) equilibration, at a reference pressure of 1.0 bar. Using periodic boundary conditions (PBC), long-range electrostatics interactions were computed using the Particle Mesh Ewald (PME) electrostatics algorithm, with a Fourier grid spacing of 1.6 Å within a cut-off radius of 12 Å in all three dimensions. This well equilibrated system was then subjected to 100 ns production of MD run utilizing the leap-frog algorithm, with trajectories generated every 10 ps with an integration time frame of 2 fs. The resulting 100 ns MD run trajectories were analyzed for the calculation of Root Mean Square Deviation (RMSD) values and the number of hydrogen bonds. All graphs were generated using XMgrace software.

### Cloning of nanobody

The gene sequence for the native nanobody was acquired from the Huo et.al 2020 ^43^. The codon-optimized sequence of the nanobody gene was synthesized by Invitrogen, Thermo Fisher Scientific. Following synthesis, the nanobody gene was PCR amplified using 5’- CAGCCATATGCAGGTTCAGCTGGTTGAA-3’ as forward and 5’- GGTGCTCGAGTTAATGATGATGGTGATGAT-3’ as reverse primer. After restriction digestion, the nanobody gene was cloned into the pET28c vector, positioned between the *NdeI* and *XhoI* restriction sites with an N-terminal histidine tag. Subsequently, the recombinant vector was transformed into competent *E. coli* DH5α cells and grown on Luria Bertani (LB) agar plates supplemented with 50 µg/mL kanamycin overnight at 37 °C. The following day, kanamycin-positive colonies were selected from the agar plate and plasmids were extracted using the MiniPrep isolation kit (Qiagen, USA). The cloned construct was eventually confirmed using Sanger sequencing.

### Site-directed mutagenesis (SDM)

After confirming the plasmid through Sanger sequencing, site-directed mutagenesis was performed to introduce selected mutations in the CDRs of the native nanobody. Specifically designed overlapping primers were used for the PCR amplification of the native nanobody gene. The amplification process individually introduced 9 single mutations, resulting in one amino acid change per mutation. Following the amplification, the parental strand was enzymatically cleaved using *DpnI*, and digested product was transformed into competent XL-1 blue cells of *E. coli* and grown on the LB agar plates containing kanamycin (50 µg/mL) and chloramphenicol (35 µg/mL). The following day, positive colonies were picked from the agar plate and the plasmids were isolated. These plasmids were sent for Sanger sequencing to confirm the presence of the intended mutations in the nanobody gene. The list of primers employed for site-directed mutagenesis are listed in (Supplementary Table 2).

### Recombinant nanobody production

The recombinant plasmid with the cloned nanobody gene was transformed into competent *E. coli* Rosetta cells (DE3; Novagen, USA) and grown on LB agar plates containing kanamycin (50 µg/mL) and chloramphenicol (35 µg/mL). For bacterial expression of nanobody protein, a single colony was picked and inoculated into 10 mL of LB broth supplemented with kanamycin (50 µg/mL) and chloramphenicol (35 µg/mL), and the culture was incubated at 37 °C and 200 rpm overnight. This primary culture was used to inoculate the 1 L LB broth, culture was grown at 37 °C till optical density (OD) OD_600_ reached 0.4. Subsequently, recombinant nanobody protein was expressed by adding 0.2 mM isopropyl-β-d-1-thiogalactopyranoside (IPTG) as inducer, and culture was grown at 16 °C for 20 h in an incubator shaker at 180 rpm. Following this, the cells were harvested by centrifugation at 4 °C at 6,000 rpm for 10 min. The cell pellet was then resuspended in the lysis buffer (50 mM Sodium Phosphate, 500 mM NaCl, and 10 mM Imidazole, pH 8.0). Cell disruption was done using a French press (Constant Systems Ltd, Daventry, England) and the cellular debris was removed by centrifugation of cell lysate at 10,000 rpm for 90 min at 4 °C. The clarified supernatant was loaded onto a pre-equilibrated gravity flow column of nickel-nitrilotriacetic acid (Ni-NTA) beads (Qiagen, Germantown, MD, USA). The column was washed with the increasing concentration of imidazole and the recombinant nanobody protein was eluted at a gradient of 250-500 mM imidazole concentration in elution buffer (50 mM Sodium Phosphate, 300 mM NaCl, and 5% glycerol, pH 8.0). Fractions collected at different imidazole concentrations were analyzed using 15% sodium dodecyl sulphate-polyacrylamide gel electrophoresis (SDS-PAGE) to confirm the purity of purified nanobody proteins (Supplementary Fig 1A). Fractions containing the pure nanobody protein were pooled together, dialyzed against 1 X phosphate-buffered saline (PBS) containing 140 mM NaCl, 2.6 mM KCl, 10 mM NaHPO_4_.2H_2_O and 1.8 mM KH_2_PO_4_ pH 7.4 with 10% glycerol for overnight at 4 °C. After dialysis the protein was concentrated through Amicon centrifugal filters (Millipore, Burlington, MA, USA) with a molecular weight cutoff of 3 kDa. The concentrated protein was flash frozen in liquid nitrogen and stored at −80 °C until further use. The expression and purification of all nanobody mutants were carried out similarly.

### Recombinant RBD production

The expression and purification for SARS-CoV-2 RBD (residues 319-528) was conducted following the previously reported protocol with some modifications ^63^. In brief, the RBD gene was cloned in pPick.9 vector in between the *AvrII* and *SnaBI* restriction sites. Upon confirming the clone through Sanger sequencing, the plasmid was linearised with *SalI* enzyme and used for transfection into competent *Pichia pastoris* cells via electroporation. The transfected cells were grown on a histidine-deficient medium for 2-3 days. Positive colonies were screened using a gentamycin gradient to identify colonies with high expression of the RBD protein. Selected colonies were subjected to a small-scale expression of RBD, and the maximum protein yielding colony was chosen. For protein expression, yeast culture was grown in buffered glycerol media medium (BMGY) at 28 °C in a shaking incubator until an OD_600_ of ∼ 2–6. The yeast cells were harvested through centrifugation and then resuspended in buffered methanol-complex medium (BMMY) containing 1% methanol, followed by growth at 28 °C for 4 days. After the 4-day induction period with methanol, the culture supernatants were collected and purified by Ni-NTA affinity chromatography using a purification buffer consisting of 50 mM Tris, 150 mM NaCl, and 10% glycerol pH 8.0. Fractions collected at different imidazole concentrations were analyzed using 12% SDS-PAGE to confirm the purity of purified RBD proteins (Supplementary Fig 1B). Fractions containing the pure RBD protein were pooled together and subsequently dialyzed against 1X PBS at pH 7.4 overnight at 4 °C. Subsequently, the protein was concentrated using Amicon centrifugal filters (Millipore, Burlington, MA, USA) with a molecular weight cut off of 10 kDa.

### Binding affinity assessment using ITC

The thermodynamic parameters and binding affinity of RBD with the nanobodies were determined through ITC with a Micro Cal ITC200 microcalorimeter (Malvern, Northampton, MA) at 25 °C. Purified RBD and native/mutant nanobody were dialyzed in 1 X PBS at pH 7.4 and 10% glycerol. The nanobody (syringe) at a concentration of 50 µM was titrated against 2.5 µM RBD (cell) ^44^. The experimental setup consists of an initial injection of 0.5 µL followed by 13 injections of 2.9 µL of the purified nanobody, into the cell containing RBD protein. The acquired data were fitted into the single-site binding model to generate the binding isotherm and calculate the dissociation constant (K_D_) for the binding. Data processing was carried out using MicroCal analysis software, Malvern, in conjunction with the commercially available Origin 7.0 program.

### Production of SARS-CoV-2 pseudovirus

Human embryonic kidney (HEK-293T) cells (NCCS, India) were used for the transient expression of SARS-CoV-2 spike pseudotyped virus. HEK-293T cells expressing Human angiotensin-converting enzyme-2 (hACE2) and TMPRSS2 were obtained from BEI Resources (NR-55293) and used for the neutralization assays. All cell lines were maintained at 37 °C and 5% CO_2_ in high glucose Dulbecco’s-modified essential media (DMEM; HiMedia, India) supplemented with 10% fetal bovine serum (FBS; Gibco, USA) and 100 U/mL penicillin and 100 mg/mL streptomycin (HiMedia, India).

The plasmid constructs for the production of pseudotyped lentiviral particles expressing S-protein of SARS-CoV-2 were obtained from the BEI Resources SARS-Related Coronavirus 2, Wuhan-Hu-1 Spike-Pseudotyped Lentiviral Kit V2 (NR-53816). Pseudovirus particles were produced following the protocol described by ^64^. Briefly, 3 × 10^6^ HEK-293T cells were seeded in 6 well plates prior to the day of transfection. At 70-80% confluency, cells were transiently transfected with different concentrations of the plasmids (NR-53742, NR-52516, NR-52517, NR-52518, and NR-52519) required to produce lentiviral particles using Lipofectamine 2000 (Invitrogen) as per the manufacturer’s instructions. After 60 h post-transfection, the culture supernatant containing pseudovirus particles was harvested and filtered using a 0.45 µm filter and stored at −80 °C in smaller aliquots. Furthermore, the virus titer was determined by luciferase assay as mentioned by Crawford et al 2020 ^64^.

### Pseudovirus neutralization assay

To perform the neutralization assay, 96 well plates were coated with 0.1 % gelatine (Himedia TCL-059). HEK-293T cells expressing human ACE2-TMPRSS2 (NR:55293) were seeded at a density of 4 ×10^4^ cells /well. On the following day, a 50 µM concentration of nanobodies were two-fold serially diluted in 2% DMEM. Serially diluted nanobodies were mixed with an equal volume of diluted virus (8×10^5^ RLU/mL) supplemented with 10 mg/mL hexadimethrine bromide (Himedia) and incubated for 1 h at 37 °C in a 5% CO_2_ incubator ^65^. Subsequently, 200 µL of each dilution was added to HEK-293T cells expressing human ACE2-TMPRSS2. Then, the plate was incubated at 37 °C in a 5% CO_2_ incubator for 60 h. Later, the cell lysate was prepared and used for luciferase assay, as mentioned in Crawford et al 2020 ^64^. From the clarified cell lysate, 20 µL of supernatant was mixed with 50 µL of luciferin substrate (SRL, India) and was added into wells of 96 well white opaque plate (Costar) to estimate the relative luminescence, using Synergy HTX multimode plate reader (Agilent BioTek). The experiment was performed thrice and data points represent the average of a triplicate set of readings. Graphs were prepared using Graph Pad Prism 8.0 software ^64,66^.

## Results

### Rationale for *in silico* mutagenesis and generation of an *in house* mutant library

CDRs of native nanobody interact with RBD and hinder the ACE2 binding and viral entry inside the cell. The native nanobody (H11-H4), originally derived from the H11 parent nanobody, was reported to be raised against RBD in Lamma ^43^. To enhance the binding affinity of native nanobody towards RBD, *in silico* affinity maturation approach was employed for which the amino acids were selected based on structural analysis of native nanobody and RBD complex generated through HADDOCK 2.4. Structural analysis of the complex revealed a major contribution of CDR2 and CDR3 loops in interactions with the RBD (Fig 1B). Furthermore, analysis using PyMol and LigPlot+ disclosed that only Arg52 and Ser57 from CDR2 were contributing in hydrogen bonding (H-bond) with the Glu484 of RBD. Whereas, CDR3 of the nanobody displayed a robust hydrogen bonding network. The residues His100, Tyr101, Val102, Tyr104, Leu106, and Asp115 of native nanobody forms H-bond with the Lys444, Asn448, Tyr449, Asn450, Glu484, Phe490, Gln493, and Ser494 residue of RBD (Fig 1C). Mutagenesis was limited to antigen recognition residues within the limit of 6 Å identified using PyMol, to minimize impact on molecular stability and enable experimental characterization with a small number of mutants. The amino acids present in the CDR2 (Arg52, Ser54, Gly55, Gly56, and, Ser57) and CDR3 (His100, Tyr101, Val102, Ser103, Tyr104, Leu105, Leu106, and Ser107) were selected and mutated using Coot software to generate an *in house* library of 112 mutants with single amino acid substitutions in the nanobody (Fig 1D). The rationale for introducing these mutations was to augment and optimize the contribution of electrostatic and polar interactions, thereby improving the overall stability and functionality through a targeted modification in the structure ^17,18^. The energy minimization step was performed using the CHARMM 27 force field in the Gromacs 2022.2 suite after the introduction of mutations in the 3D structure of the nanobody. The comparative structural alignment of the CDR loops of native/mutant nanobodies revealed that mutational changes in CDR3 not only affected CDR3 but also CDR1 and CDR2. These slight structural changes in CDRs, presumably affect the binding pattern at the interface of nanobody and RBD, subsequently increasing and decreasing the binding affinities (Supplementary Fig 3A).

**Fig. 1.**
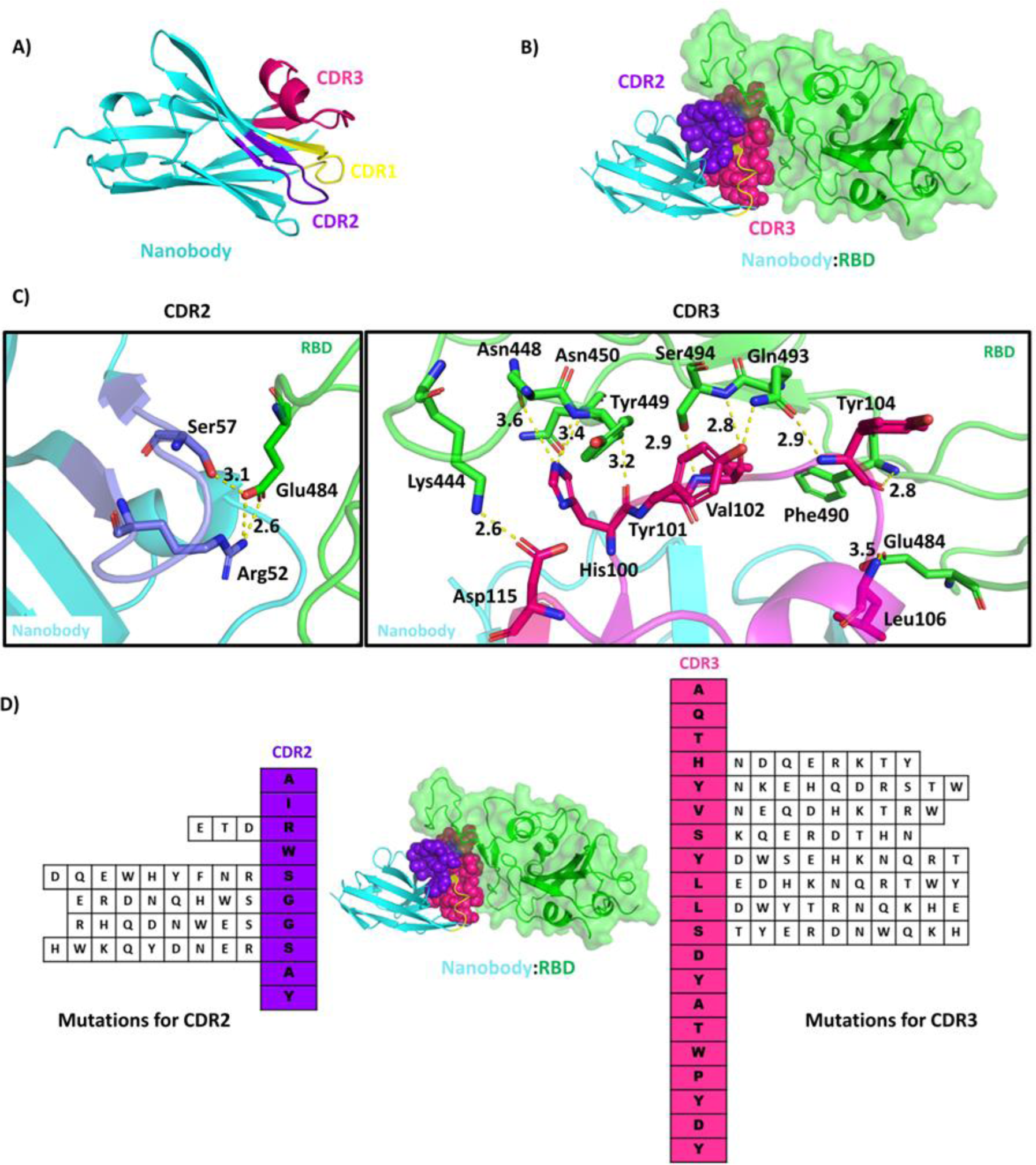
Detailed structural analysis of nanobody and RBD complex and selection of residue for the *in silico* mutagenesis. A) Cartoon representation of nanobody showing framework region (cyan) and position of CDR1 (yellow), CDR2 (purple), and CDR3 (hot pink). B) Surface representation of RBD (green) and nanobody (cyan) complex generated through HADDOCK 2.4 showing the interacting interface. The CDR2 (purple) and CDR3 (hot pink) regions of nanobody interacting with RBD are depicted as spheres. C) The key interactions present at the interface of RBD (green) and nanobody (cyan) complex generated through HADDOCK 2.4 are depicted, highlighting the CDR2 (purple) and CDR3 (magenta) of nanobody. Interacting residues of nanobody and RBD are shown in sticks. D) Vertical representation depicts the nanobody residues present in the CDR2 (purple) and CDR3 (hot Pink). The residues selected for *in silico* mutagenesis and library generation from the CDR2 and CDR3 are depicted in horizontal rows (112 mutants).

### Selection of the best binders using protein-protein docking and MD simulation

High-throughput protein-protein docking studies screened and identified the best binders compared to the native nanobody, from an *in house* library of 112 mutants generated against the RBD (Supplementary Table 1) using Coot. Additionally, a comparative analysis of interactions between native/mutated nanobody and RBD docked complexes revealed the formation of new interactions owing to the introduction of certain mutations (Fig 2 and Supplementary Fig 2). Based on interaction examination, out of the 112 mutants studied, 51 showed docking scores comparable to or higher than the native nanobody and formed additional interactions with the RBD, 9 mutants, which had not been previously published were chosen for further characterization. The mutants R52D, S54E, G56H, and S57D were selected for engineering the CDR2 loop, and L105E, L106T, L106Q, S107Q, and S107R were selected from CDR3 for further studies (Table 1). Interestingly, mutation R52D in native nanobody led to the disruption of the salt bridge between Arg52 and Glu484, rather a new H-bond was formed between Asp52 and Gln493 of RBD. For the Ser54 mutations to Glu54, the mutated residue was interacting with the RBD through weak Vander Waal interactions only. The mutation of Gly56 to His56 has imparted changes in the CDR2 loop and pulled Gly55 towards the Gly482 of RBD as observed in PyMol analysis. In native nanobody, Ser57 forms a H-bond with Glu484 of RBD and after mutation of Ser57 to Asp57, this H-bond remained intact. For the CDR3, the mutation of Leu105 to Glu105 resulted in the formation of additional polar interactions with RBD compared to native nanobody. Mutations of Leu106 to Thr106 and Gln106 resulted in polar contacts with the Glu484, the same as of native nanobody and RBD complex. The mutations of Ser107 residue to Arg107 and Gln107 resulted in an additional H-bond with Arg107 and Gly485 of RBD compared to the native nanobody and RBD complex (Fig 2 and Supplementary Fig 2).

**Fig. 2.**
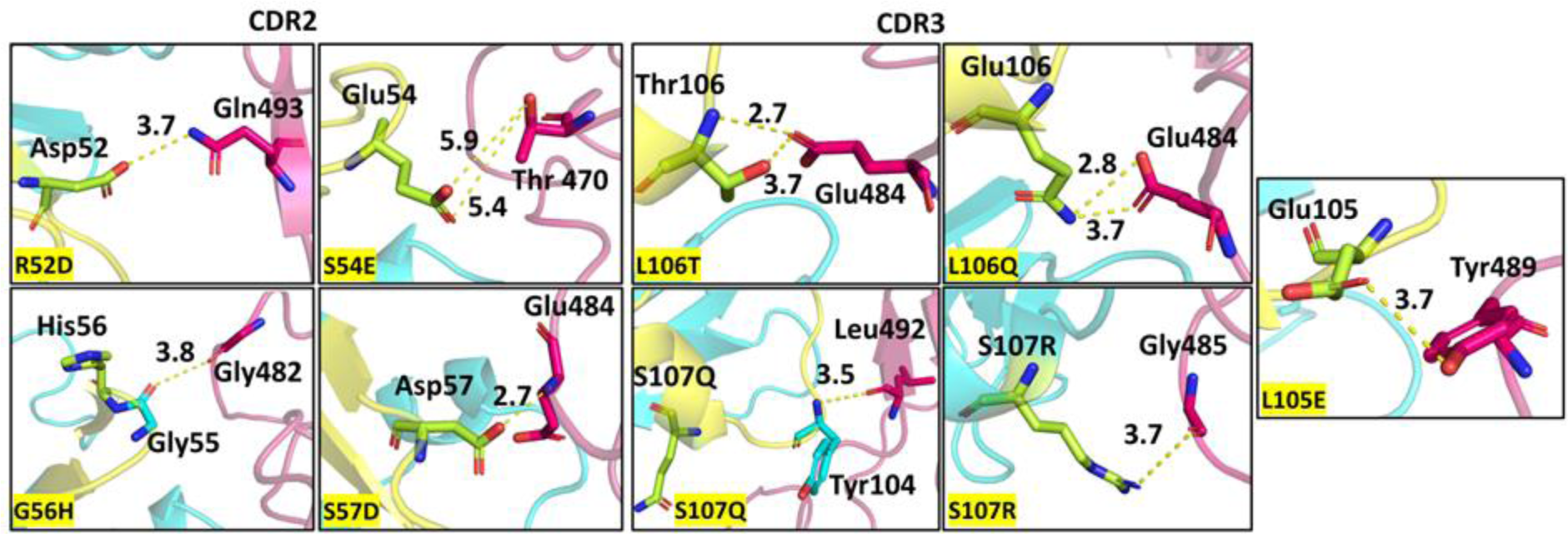
Protein-protein docking studies reveal differences in interactions of native and mutant nanobodies and RBD complexes. A comprehensive analysis was conducted on mutants of CDR2 and CDR3 of the nanobody in complex with the RBD of SARS-CoV-2, generated using HADDOCK 2.4. The cartoon representation depicts the key interacting residues of a nanobody (cyan) and RBD (light magenta) that were critically studied to understand the effects of the mutations (limon) on the binding affinity. Residues were shown as sticks using PyMol.

**Table 1.** Mutations selected for the CDR2 and CDR3 with their docking scores and number of H-bonds based on DimPlot analysis of docked complexes generated through HADDOCK 2.4 and average number of H-bonds calculated after 100 ns MD simulations run.

| **Mutation** | **HADDOCK score** | **No. of H-bond after Docking** | **No. of average H-bond after MD** |
| --- | --- | --- | --- |
| **Native nanobody** | -113.1+/-4.6 | 9 | 6.26 |
| **R52D** | -102.3+/-6.5 | 5 | 4.81 |
| **S54E** | -121.5+/-3.9 | 14 | 7.98 |
| **G56H** | -114.9+/-3.0 | 9 | 8.37 |
| **S57D** | -108.6+/-1.7 | 10 | 6.71 |
| **L105E** | -120.0+/-1.6 | 13 | 7.84 |
| **L106T** | -111.7+/-1.0 | 11 | 7.07 |
| **L106Q** | -111.8+/-2.2 | 13 | 7.60 |
| **S107Q** | -119.0+/-3.1 | 10 | 6.98 |
| **S107R** | -120.0+/-3.0 | 10 | 7.45 |

Further, for the comparative assessment of the stability and interaction pattern of these selected complexes at the atomistic level, the complexes were subjected to MD simulation studies using a CHARMM 27 ^67^ force field in the Gromacs 2022.2 suite. The average RMSD of the C_α_ backbone of complexes showed no major fluctuations, the overall values were observed to be below 0.5 Å for all complexes. Low RMSD values suggest the preservation of structural integrity and stability of complexes despite mutational changes in nanobody (Supplementary Fig 3B). The average number of H-bonds formed between RBD and native/mutant nanobody were ∼ 6-7 throughout the MD run. For some mutants, the number of H-bonds reached up to ∼ 9-10 indicating the formation of new interactions owing to the nature of mutations (Supplementary Fig 3B and Table 1). MD simulation studies indicate rearrangement of the nanobody-RBD interface residues as a result of amino acid changes present in nanobody CDRs (Supplementary Fig 4).

Docked complexes of native/mutant nanobody-RBD were analyzed to see the distribution of RBD residue at the interface of the complex within a 4 Å cutoff. All mutants were interacting with the almost same residue stretch on the RBD as the native nanobody. Here, the mutants R52D, G56H, S107Q, and S107R were among the mutants that were targeting some additional residue of RBD as shown in Fig 3. RBD and ACE2 interaction is critical for the life cycle of SARS-CoV-2. Mutations in nanobody disrupting the interaction at the ACE2-RBD interface could operate as promising entry inhibitors for SARS-CoV-2. Interestingly, analysis of RBD and native/mutant nanobody complex revealed that the RBD residues targeted by nanobody were also involved in interaction with ACE2 (Fig 3 and Supplementary Table 3). Additionally, a few mutations in nanobody were identified to target some additional residue of the ACE2 binding pocket of RBD (Fig 3 and Supplementary Table 3). These mutant nanobodies presumably act like molecular roadblocks, preventing virus entry and infection.

**Fig. 3.**
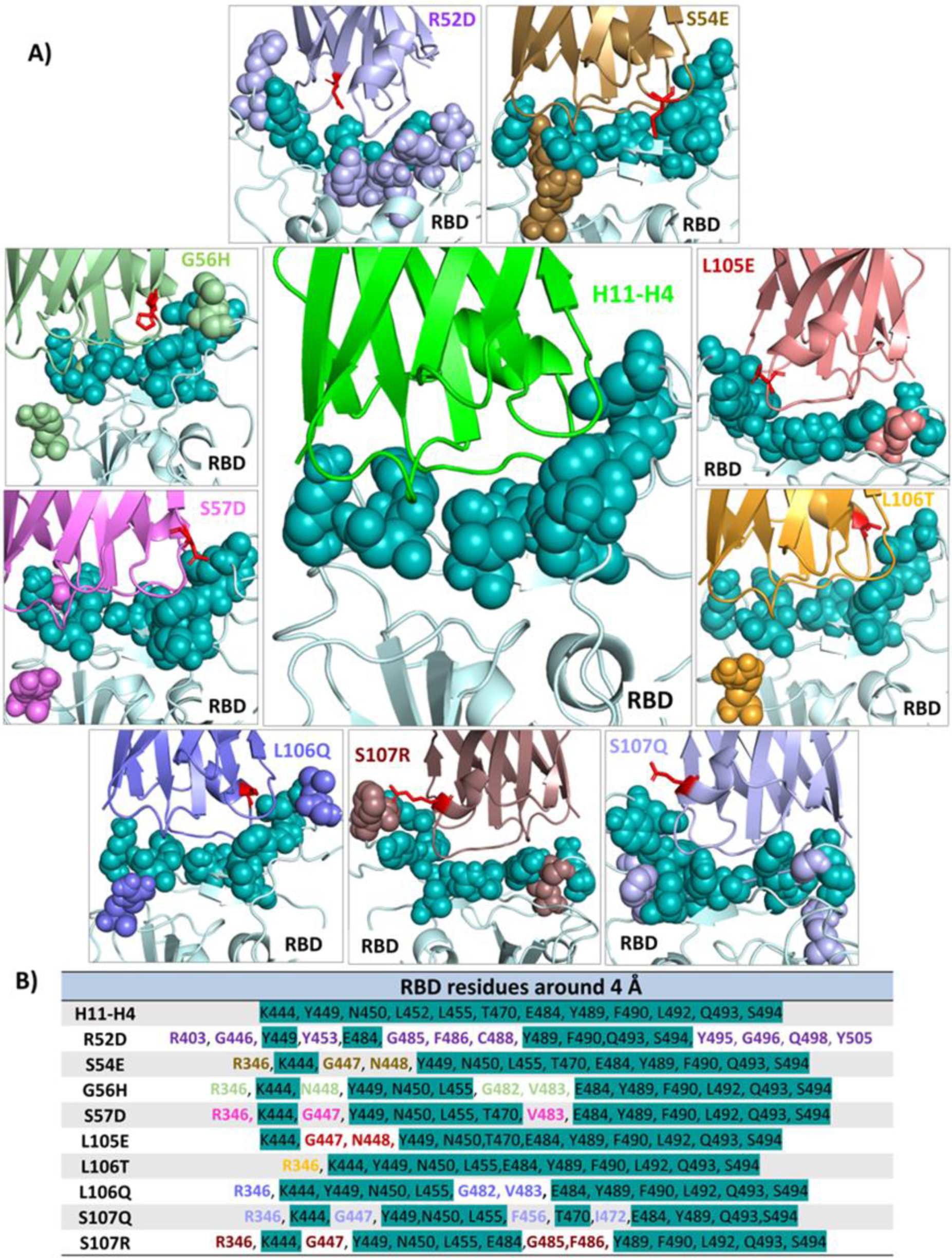
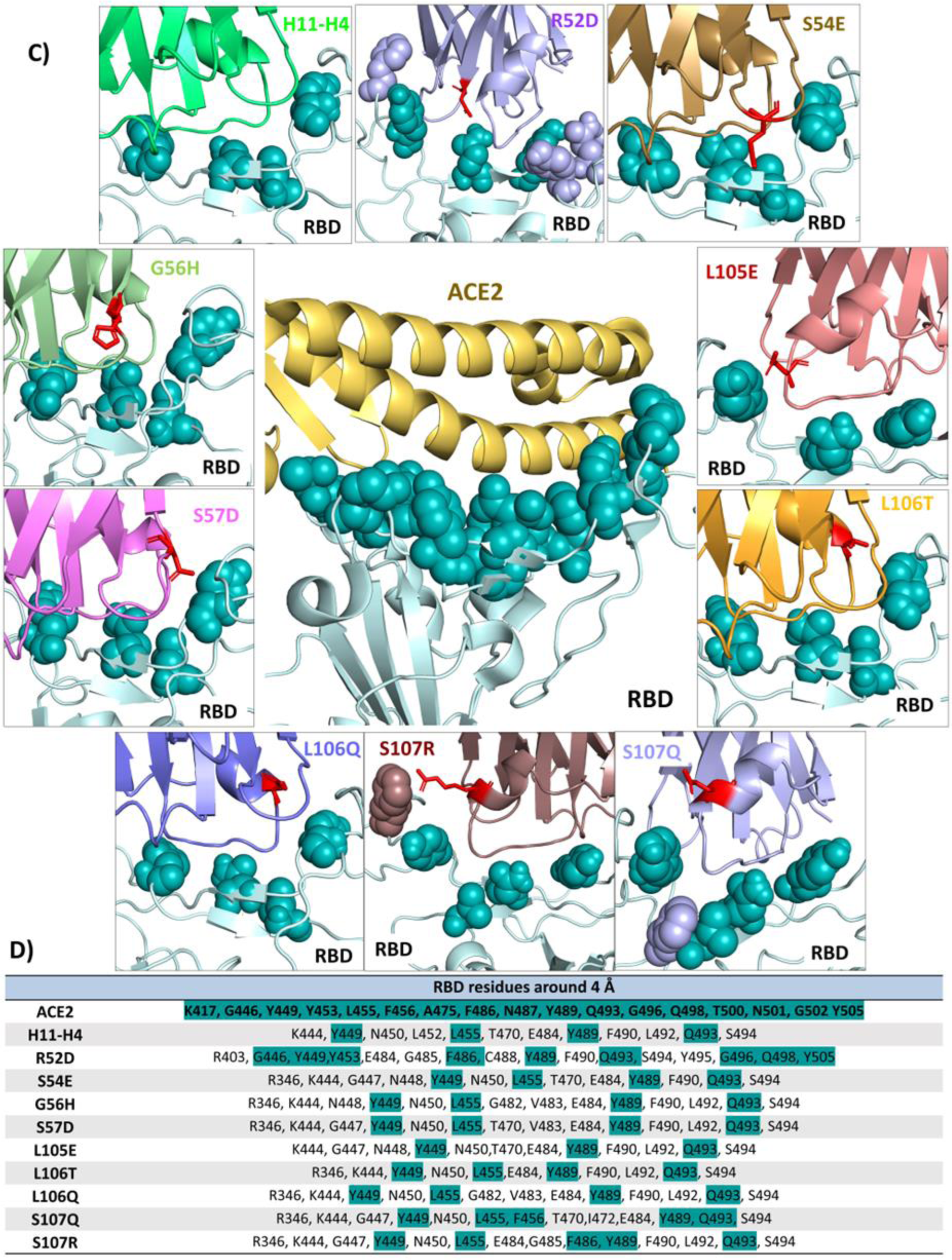
The comparative analysis of interacting residues at the interface of native and mutant nanobody-RBD complex and ACE2-RBD (PDB: 6MOJ) complex. A) In the native/mutant nanobody-RBD (pale cyan) docked complexes, RBD residues within 4 Å cutoff were shown in teal spheres. For the mutants of nanobody, RBD residues were colour coded in two formats: teal if residues same as native nanobody and in mutation-specific colors. Mutated residue in nanobody were shown in red sticks. B) Tabular representation for RBD residues in a colour coded manner. For the native/mutant nanobody-RBD complex, residues targeted by native nanobody were highlighted in cyan, additional RBD residues targeted by mutants were colour coded as per mutation. C) In the RBD (pale cyan)-ACE2 (yellow orange) complex, RBD residues within the 4 Å cutoff were shown in teal sphere. In the docked complexes of native/mutant nanobody-RBD residues same as of ACE2-RBD complex were shown in teal spheres, while other additional RBD residue were shown in mutation-specific colors. Mutated residue in nanobody were shown in red sticks. D) Tabular representation of the RBD residues within the 4 Å cutoff of ACE2-RBD complex and native/mutant nanobody-RBD complex, highlighting overlapping residues in cyan. Figure prepared using PyMol.

### Validation of computational predictions by assessment of binding affinities through ITC

To establish the accuracy of the *in silico* findings, binding affinities of the mutants with the SARS-CoV-2 RBD were estimated using ITC. The binding titrations of the RBD with the nanobody are exothermically driven. Thermodynamic data generated from ITC isotherms was fitted into a single-site binding model to compare the binding affinities of the native and mutants. Thermodynamic parameters and titration curve depicted that the introduction of mutations such as S54E, S57D, L105E, L106T, L106Q, S107R, and S107Q contributed to the improved affinities towards the RBD, compared to the native nanobody (Fig 4). Notably, substitutions involving charged or polar residues tend to be particularly effective in improving binding affinity, highlighting the significance of these interactions in computational affinity enhancement, which is consistent with *in silico* findings.

**Fig. 4.**
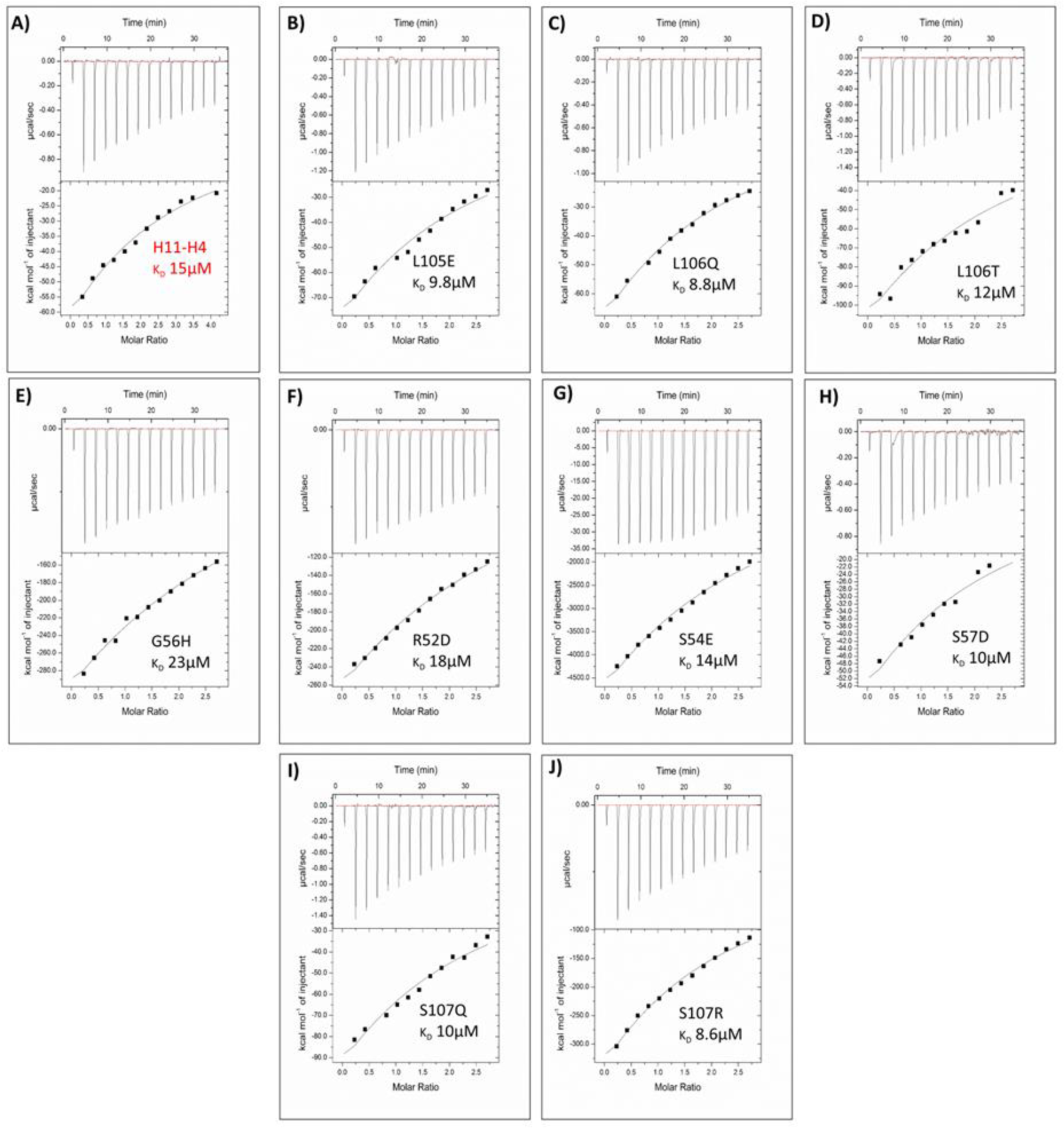
Determination of binding affinity of native /mutant nanobody with RBD using ITC. (A-J) Binding isotherm plots were generated using a single-site binding model with the binding affinities of the native/mutants. The upper portion of each ITC titration depicts raw data, while the lower portion displays the fitted binding isotherm, with inset K_D_ values provided for the native/mutants.

### Potent pseudovirus neutralization by generated mutant nanobodies

The replication of pseudotyped lentiviral particles expressing S-protein successfully mimics SARS-CoV-2 host cell entry ^64^. Consequently, pseudotyped lentiviral particles expressing surface exposed S-protein of SARS-CoV-2 were produced and used to determine the neutralization efficacy of native and the mutant nanobody using HEK-293T cells expressing human ACE2-TMPRSS2. Pseudotyped lentiviral particles were pretreated with increased concentrations of nanobody, and incubated with HEK-293T cells expressing human ACE2-TMPRSS2 for 60 h. The cell lysate was prepared following the method described previously ^64^. The supernatant (20 µL) from clarified cell lysate was mixed with luciferin substrate (50 µL/well) in a white opaque 96-well plate. Relative luminescence was measured using a Synergy HTX multimode plate reader (Agilent BioTek). The IC50 values were computed using Graph Pad Prism 8.0 software. The experiment was conducted three times, and each data point represents the average of three readings. Neutralizing efficiency varied considerably among the different mutants with IC_50_ values ranging from 0.03 −15.27 µM. Interestingly the mutants L106T, L106Q, S107R, and S107Q have neutralization efficiency significantly better than the native nanobody which is in concordance with the biophysical studies (Fig 5).

**Fig. 5.**
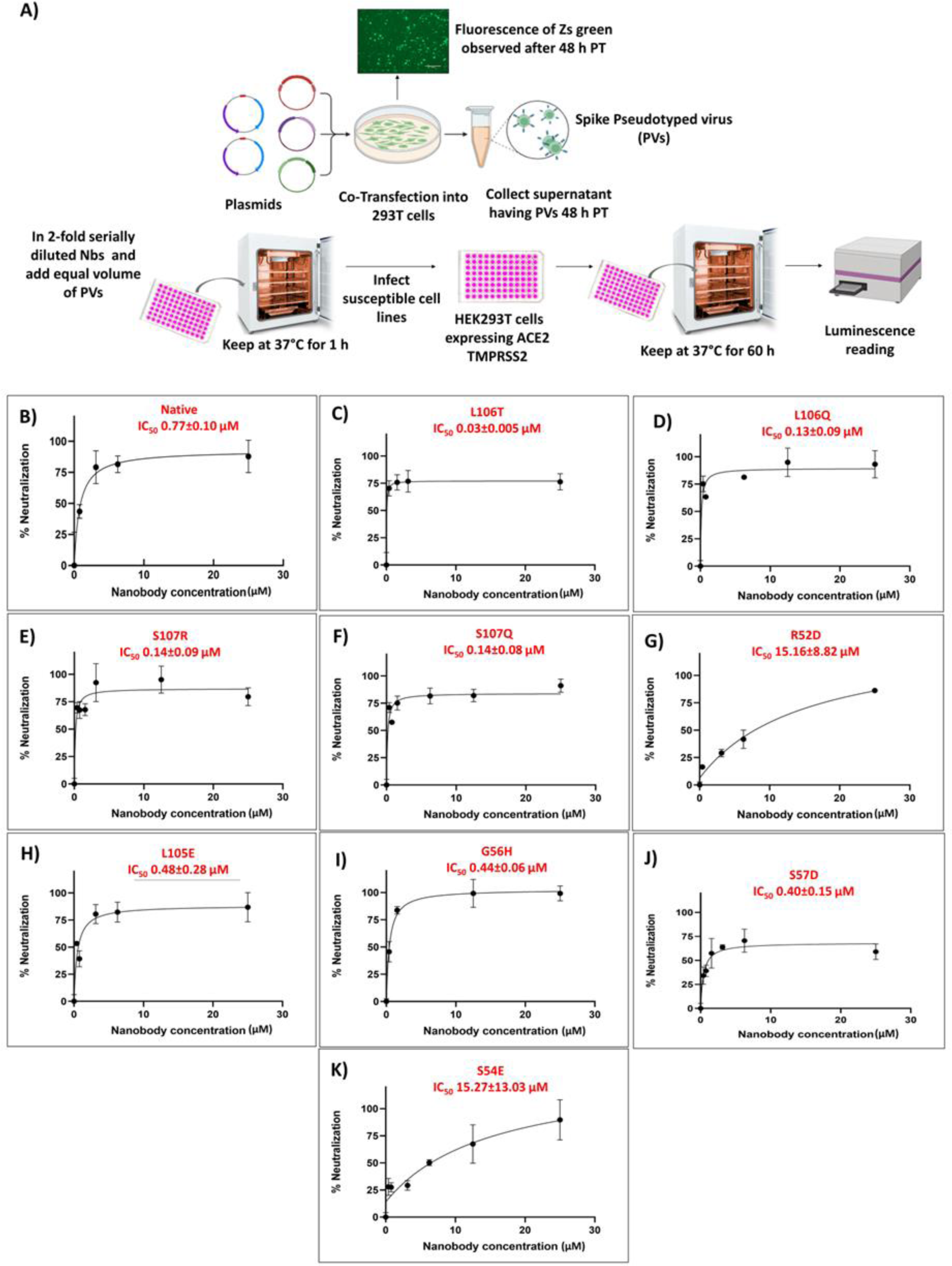
Evaluation of neutralization efficiency of native and mutant nanobody using pseudovirus assay in HEK-293T cells expressing human ACE2-TMPRSS2. A) Schematic for the production of SARS-CoV-2 pseudovirus and neutralization assay. (B-K) The neutralization potency of nanobodies were determined using a luciferase based pseudotyped SARS-CoV-2 neutralization assay. Pseudotyped lentiviral particles expressing S-protein of SARS-CoV-2 were pretreated with increased concentrations of nanobody and subsequently incubated with HEK-293T cells expressing human ACE2-TMPRSS2 for 60 h. Post infection, luciferase activity in cell lysates was quantified to calculate IC_50_ values. The IC_50_ values were computed using Graph Pad Prism 8.0 software. The experiment was conducted three times, and each data point represents the average of three readings.

## Discussion

The COVID-19 pandemic has impacted the world’s population by disrupting social, mental, and economic harmony. The year-long vaccination drive against SARS-CoV-2 has yielded positive results, but the efficacy of current SARS-CoV-2 vaccines and treatments is increasingly challenged by the virus’s capacity to evolve. To keep pace with viral evolution, a fast, potent, and cost-effective preventative or therapeutic strategy is the need of the hour. While vaccines prime our immune system to create antibodies against the virus, antibodies-based therapies offer a different approach. They can be directly administered to patients, providing immediate protection or treatment, particularly for those with weakened immune systems who may not respond well to vaccines ^5^.

The success of Caplacizumab, the first FDA-approved nanobody for acquired thrombotic thrombocytopenic purpura (aTTP) treatment, highlights the therapeutic potential of nanobodies. This is further supported by promising results from animal studies using nanobodies as preventative measures (prophylactics) ^68–70^. Owing to their small molecular weight, the nanobodies can engage relatively well with large antigenic interfaces. Nanobodies are versatile and easy to manipulate hence serve as good alternatives to conventional antibodies ^26^. In the last two years, several single-domain antibodies i.e. nanobodies have been isolated from the naïve library that binds to the RBD of SARS-CoV-2 ^41,43,44^. Nanobodies can be affinity matured for improved binding to target antigens using both experimental and computational tools. *In silico* affinity maturation is robust, efficient, and less laborious compared to conventional methods such as phage display, yeast display, error-prone PCR (Polymerase Chain Reaction), and DNA Shuffling ^22,71^. Mutating the residues of paratopes (antigen-binding site) at the interface of the antigen-antibody complex can significantly impact the complex’s affinity. *In silico* affinity maturation strategies often involve structure-guided mutations and the calculation of changes in thermodynamic parameters resulting from these mutations ^21,43,72^. This approach of *in silico* affinity maturation has been successfully applied to antibodies and antibody fragments such as Fab, scFvs, and nanobodies ^17,21,22,71,73,74^.

For an antigen antibody complex, around eighty percent of binding energy is reported to be contributed by a small number of significant interactions, which are predominantly found in the CDRs ^17^. In earlier reports, methods based on computational prediction such as the ADAPT (Assisted Design of Antibody and Protein Therapeutics) platform, identified 20 to 30 targeted mutations per system to achieve the desired affinity enhancements ^73^. In this study, structure-guided *in silico* mutagenesis pipeline was used for the affinity maturation of the native nanobody, which binds to the RBD of SARS-CoV-2 and hinders the interaction of RBD with ACE2. Detailed structural analysis of the native nanobody and RBD complex revealed the hotspot residue within CDR2 and CDR3, that were further selected for *in silico* mutagenesis and docking studies (Fig 1B). A structure-assisted virtual library of 112 mutants was generated and screened against RBD (Supplementary Table 1). Of the 112 mutants, 9 single mutations were shortlisted, based on docking score, favourable interactions, and MD simulations studies (Table 1). To validate the findings of *in silico* work, the native /mutates were produced recombinantly used for the *in vitro* evaluation of binding affinities using ITC. In the mutations of CDR2 and CDR3, the formation of additional interactions would have contributed in enhancing the binding affinity towards RBD (Fig 2 and Fig 4). Further, the lentiviral-based pseudovirus neutralization assay identified four potent virus-neutralizing mutants: L106T, L106Q, S107R, and S107Q (0.03 µM, 0.13 µM, 0.14 µM and 0.14 µM respectively) having improved neutralization potential, compared to the native nanobody (IC_50_ = 0.77 µM) (Fig 5).

In previous reports of the native nanobody Arg52 is reported to form a bivalent salt linkage with Glu484. Additionally, this Arg52 is also documented to form a H-bond with the main-chain Ser103 and Tyr109 ^43^. Substituting Arg52 with Glu52 disrupted the salt linkage and eventually diminished the binding affinity towards RBD, as reported ^75^. Owing to the imperative role of Arg52 in nanobody-RBD interaction, the residue was selected and further subjected to the developed *in silico* pipeline. Intriguingly, mutating Arg52 to Asp52 in the present study resulted in disruption of the salt-bridge as indicated *in silico* studies. In concordance with the *in silico* findings, a decreased binding affinity and neutralization potency was observed for the R52D mutant in comparison to the native nanobody, thereby validating the applicability of the developed pipeline (Fig 5). Across *in silico* and *in vitro* investigations, the mutations most frequently observed in this study entail introducing charged or polar residues into the sequence, which emphasizes the role of these residues in affinity enhancement as also reported earlier ^21,22,76,77^.

Published reports describe the development of optimized antibodies with multiple mutations, ranging from three to fourteen, in their primary sequences, without utilizing electrostatic optimization ^71,78^. In contrast, our approach achieved a 1.5-fold increase in affinity for native nanobody with just a single mutation, which is less likely to alter the 3D structure of nanobody.

In summary, four potential mutants of native H11-H4 nanobody with enhanced affinity against SARS-CoV-2 RBD have been generated in this study using structure-guided *in silico* approach, which is cost effective and less time consuming. Mutations in non-active site residues can significantly impact distant sites, altering substrate specificity through cooperative effects this is well-documented and frequently observed in directed evolution experiments ^79^. Mutations in CDR2 and CDR3 might have induced structural changes in CDR1 or the framework region of the nanobody, affecting its affinity and neutralization efficacy upon mutation. Follow up *in vitro* virus-based, *in vivo* mouse model studies and determination of the atomic structure of Ag-Ab complex (RBD and mutant) is next step for the study and will help to unravel the underlying physicochemical inhibitory mechanisms of engineered nanobodies in detail. The developed pipeline can be used easily and effectively to generate CDRs tailored to emerging variants of SARS-CoV-2 in the near future.

## Conclusion

The present study illustrates the potential of computational techniques in refining antibodies for enhanced affinity towards specific antigens, as evidenced by the *in silico* affinity maturation of a native nanobody targeting the RBD of SARS-CoV-2. Identified key mutations that significantly improved the binding affinity showcased the effectiveness of the designed structure-guided *in silico* mutagenesis pipeline. The findings underscore the importance of computational methods in antibody optimization, complemented by rigorous biophysical characterization and *in vitro* efficacy validation. This integrated approach holds promise for accelerating the development of therapeutic interventions against evolving pathogens like SARS-CoV-2 and others.

## Acknowledgment

ST and JS acknowledge and thank the Indian Council for Medical Research (ICMR) Government of India (Project. ref no ISRM/12/ (06)/2022) for supporting this study. VS is thankful to ICMR and MB, SC are thankful to the Council of Scientific and Industrial Research (CSIR) for financial support. SKN acknowledges the Ministry of Human Resource Development, (MHRD) for research fellowship. ST, JS and PK thanks the Department of Biotechnology, Govt of India for supporting the Translational and Structural Bioinformatics Centre at Department of Biosciences and Bioengineering, IIT Roorkee (reference number BT/PR40141/BTIS/137/16/2021). Authors also thank Ashok Soota Molecular Medicine Facility and Macromolecular Crystallographic Unit (MCU) at the Indian Institute of Technology Roorkee (IIT Roorkee).

## Authors contributions

Conceptualization, S.T., J.S., R.K., and P.K.; methodology, S.T., V.S., M.B., and S.C.; experimentation, V.S., M.B., S.C., and S.K.N.; formal analysis, S.T., J.S., R.K., P.K., VS., M.B., and S.C.; writing-original draft, S.T., V.S., M.B., and S.C., writing-review & editing, S.T., J.S., R.K., and P.K. supervision, S.T., and P.K.; All authors read, revised and approved the manuscript.

## Supplementary Tables

**Supplementary Table 1:** The docking score, number of salt bridge and hydrogen bond contacts calculated using Dimplot for the 112 mutations generated using Coot on CDR2 and CDR3 loops of native nanobody. Mutants selected for further MD simulation studies are highlighted in light green.

|  | Mutation | HADDOCK score | Number of Hydrogen bonds | Number of Salt bridge |
| --- | --- | --- | --- | --- |
|  | Native nanobody (Reference) | -113.1+/-4.6 | 9 | 0 |
| 1. | R52D | -102.3+/-6.5 | 5 | 0 |
| 2. | R52E | -103.5+/-7.4 | 6 | 0 |
| 3. | R52S | -101.4+/-1.8 | 5 | 2 |
| 4. | S54R | -119.7+/-3.7 | 10 | 0 |
| 5. | S54N | -116.9+/-1.2 | 11 | 1 |
| 6. | S54D | -118.3+/-2.0 | 10 | 0 |
| 7. | S54Q | -117.0+/-2.6 | 9 | 0 |
| 8. | S54E | -121.5+/-3.9 | 14 | 0 |
| 9. | S54H | -118.1+/-1.6 | 12 | 0 |
| 10. | S54F | -116.4+/-0.8 | 8 | 1 |
| 11. | S54Y | -125.3+/-2.8 | 9 | 0 |
| 12. | S54W | -126.1+/-1.5 | 11 | 0 |
| 13. | G55R | -113.7+/-5.1 | 6 | 0 |
| 14. | G55S | -112.9+/-1.0 | 8 | 1 |
| 15. | G55H | -122.7+/-2.0 | 10 | 0 |
| 16. | G55E | -98.5+/-7.2 | 9 | 0 |
| 17. | G55Q | -110.5+/-1.1 | 11 | 0 |
| 18. | G55D | -113.6+/-1.7 | 10 | 0 |
| 19. | G55N | -118.1+/-1.6 | 12 | 0 |
| 20. | G55W | -128.7+/-2.3 | 10 | 0 |
| 21. | G56N | -112.1+/-2.6 | 8 | 0 |
| 22. | G56Q | -116.9+/-1.5 | 11 | 0 |
| 23. | G56R | -113.6+/-1.5 | 9 | 0 |
| 24. | G56D | -111.0+/-2.3 | 10 | 0 |
| 25. | G56E | -92.8+/-5.0 | 3 | 0 |
| 26. | G56H | -114.9+/-3.0 | 9 | 0 |
| 27. | G56S | -113.2+/-1.7 | 6 | 1 |
| 28. | G56W | -114.7+/-1.5 | 9 | 0 |
| 29. | S57R | -108.9+/-7.9 | 11 | 0 |
| 30. | S57Y | -121.5+/-3.8 | 8 | 1 |
| 31. | S57W | -114.8+/-3.2 | 8 | 0 |
| 32. | S57K | -119.4+/-4.3 | 11 | 1 |
| 33. | S57H | -125.9+/-2.1 | 10 | 0 |
| 34. | S57E | -95.8+/-1.8 | 5 | 0 |
| 35. | S57Q | -92.0+/-1.8 | 4 | 0 |
| 36. | S57D | -108.6+/-1.7 | 10 | 0 |
| 37. | S57N | -118.3+/-4.7 | 12 | 0 |
| 38. | H100D | -112.9+/-3.4 | 8 | 0 |
| 39. | H100E | -117.7+/-2.6 | 7 | 1 |
| 40. | H100N | -104.1+/-1.5 | 9 | 0 |
| 41. | H100K | -107+/-1.7 | 8 | 0 |
| 42. | H100Y | -111.1+/-0.7 | 10 | 0 |
| 43. | H100T | -109.3+/-0.7 | 11 | 0 |
| 44. | H100Q | -113.2+/-2.7 | 9 | 1 |
| 45. | H100R | -111.9+/-0.1 | 10 | 0 |
| 46. | Y101K | -109.9+/-1.3 | 8 | 1 |
| 47. | Y101N | -112.2+/-0.3 | 7 | 0 |
| 48. | Y101S | -110.9+/-3.0 | 9 | 1 |
| 49. | Y101E | -108.3+/-4.9 | 10 | 0 |
| 50. | Y101D | -95.3+/-3.5 | 10 | 0 |
| 51. | Y101W | -100.7+/-2.0 | 3 | 0 |
| 52. | Y101R | -105.5+/-2.8 | 11 | 0 |
| 53. | Y101Q | -99.2+/-1.3 | 7 | 0 |
| 54. | Y101H | -97.5+/-3.9 | 7 | 0 |
| 55. | Y101T | -120+/-3.5 | 10 | 0 |
| 56. | V102W | -120.7+/-1.3 | 7 | 1 |
| 57. | V102R | -108.6+/-5.5 | 7 | 0 |
| 58. | V102Q | -105.5+/-2.4 | 8 | 0 |
| 59. | V102K | -109.2+/-2.9 | 10 | 1 |
| 60. | V102N | -97.6+/-1.1 | 5 | 0 |
| 61. | V102H | -116.3+/-3.7 | 10 | 0 |
| 62. | V102E | -96.8+/-2.8 | 6 | 0 |
| 63. | V102D | -110.7+/-1.1 | 9 | 1 |
| 64. | V102T | -109.4+/-1.3 | 7 | 1 |
| 65. | S103E | -122.1+/-3.7 | 11 | 1 |
| 66. | S103R | -122.1+/-1.8 | 11 | 0 |
| 67. | S103T | -123.5+/-1.3 | 12 | 0 |
| 68. | S103D | -117.2+/-3.7 | 11 | 0 |
| 69. | S103H | -116.5+/-2.5 | 9 | 0 |
| 70. | S103N | -113.8+/-1.8 | 10 | 1 |
| 71. | S103Q | -119.3+/-1.0 | 12 | 0 |
| 72. | S103K | -114.2+/-2.0 | 5 | 1 |
| 73. | Y104E | -91.7+/-2.0 | 4 | 1 |
| 74. | Y104H | -96.8+/-4.7 | 7 | 0 |
| 75. | Y104K | -95.9+/-6.3 | 4 | 1 |
| 76. | Y104N | -83.4+/-4.0 | 6 | 0 |
| 77. | Y104S | -106.5+/-0.7 | 13 | 0 |
| 78. | Y104Q | -91.8+/-6.2 | 11 | 1 |
| 79. | Y104R | -88.4+/-5.7 | 6 | 1 |
| 80. | Y104D | -91.3 +/- 3.2 | 5 | 2 |
| 81. | Y104W | -107.6+/-1.3 | 5 | 0 |
| 82. | Y104T | -107.0 +/- 2.8 | 12 | 0 |
| 83. | L105D | -113.7+/-2.3 | 11 | 0 |
| 84. | L105E | -120.0+/-1.6 | 13 | 0 |
| 85. | L105R | -113.2+/-2.8 | 9 | 0 |
| 86. | L105N | -115.8+/-3.7 | 12 | 0 |
| 87. | L105H | -119.1+/-6.3 | 11 | 0 |
| 88. | L105Q | -114.5+/-2.3 | 10 | 1 |
| 89. | L105K | -115.2+/-3.3 | 10 | 0 |
| 90. | L105T | -115.0+/-1.6 | 12 | 1 |
| 91. | L105Y | -125.2+/-3.2 | 10 | 0 |
| 92. | L105W | -133.3+/-1.7 | 10 | 1 |
| 93. | L106W | -106.6+/-2.7 | 5 | 1 |
| 94. | L106Y | -94.8+/-4.0 | 6 | 0 |
| 95. | L106T | -111.7+/-1.0 | 11 | 0 |
| 96. | L106R | -90.5+/-3.4 | 3 | 1 |
| 97. | L106Q | -111.8+/-2.2 | 13 | 0 |
| 98. | L106N | -114.8+/-4.0 | 11 | 0 |
| 99. | L106K | -114.7 +/- 6.6 | 11 | 0 |
| 100. | L106H | -111.9+/-6.0 | 9 | 1 |
| 101. | L106E | -94.6+/-3.8 | 8 | 0 |
| 102. | L106D | -90.8+/-3.7 | 5 | 0 |
| 103. | S107E | -117.1+/-1.5 | 8 | 1 |
| 104. | S107T | -117.3+/-3.0 | 11 | 0 |
| 105. | S107Y | -125.2+/-3.1 | 12 | 0 |
| 106. | S107N | -118.2+/-1.8 | 9 | 0 |
| 107. | S107Q | -119.0+/-3.1 | 10 | 0 |
| 108. | S107K | -117.5+/-2.0 | 9 | 0 |
| 109. | S107H | -121.0+/-1.0 | 10 | 1 |
| 110. | S107R | -120.0+/-3.0 | 10 | 0 |
| 111. | S107W | -128.6+/-4.6 | 10 | 0 |
| 112. | S107D | -114.8 +/- 2.1 | 8 | 0 |

**Supplementary Table 2:**
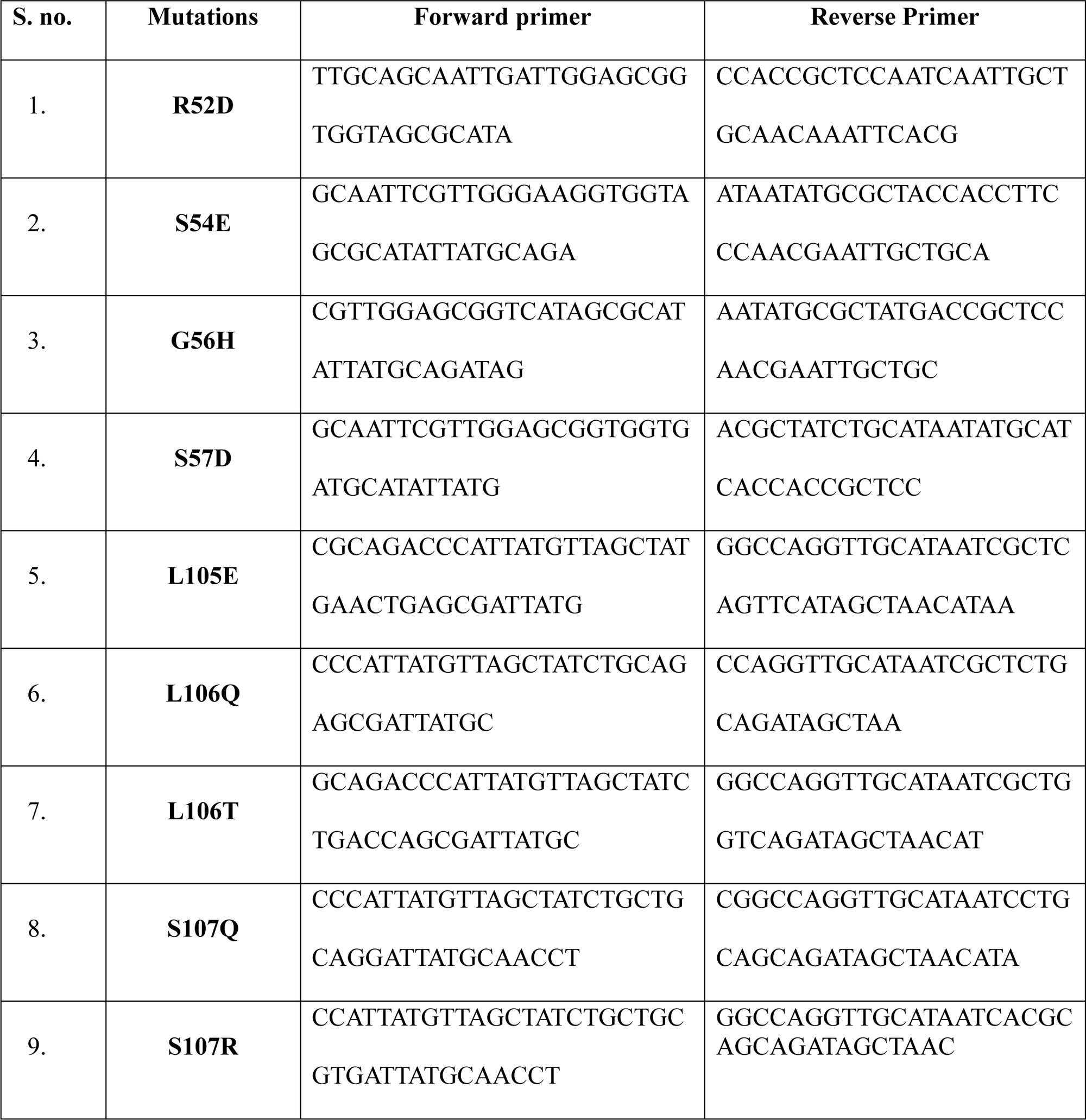
Set of primers used for the Site Directed Mutagenesis of native nanobody.

**Supplementary Table 3:** The distribution of RBD residues within 4 Å cutoff of native/mutant nanobody-RBD docked complexes and ACE2-RBD (PDB: 6MOJ) complex in colour coded manner. Here boxes adjacent to RBD residue were coloured light blue if they are present in indicated complex otherwise left blank.

| RBD residues around 4 Å |  |
| --- | --- |
| ACE2 | K417, G446, Y449, Y453, L455, F456, A475, F486, N487, Y489, Q493, G496, Q498, T500, N501, G502, Y505 |
| H11-H4 | K444, Y449, N450, L452, L455, T470, E484, Y489, F490, L492, Q493, S494 |
| R52D | R403, G446, Y449, Y453, E484, G485, F486, C488, Y489, F490, Q493, S494, Y495, G496, Q498, Y505 |
| S54E | R346, K444, G447, N448, Y449, N450, L455, T470, E484, Y489, F490, Q493, S494 |
| G56H | R346, K444, N448, Y449, N450, L455, G482, V483, E484, Y489, F490, L492, Q493, S494 |
| S57D | R346, K444, G447, Y449, N450, L455, T470, V483, E484, Y489, F490, L492, Q493, S494 |
| L105E | K444, G447, N448, Y449, N450, T470, E484, Y489, F490, L492, Q493, S494 |
| L106T | R346, K444, Y449, N450, L455, E484, Y489, F490, L492, Q493, S494 |
| L106Q | R346, K444, Y449, N450, L455, G482, V483, E484, Y489, F490, L492, Q493, S494 |
| S107Q | R346, K444, G447, Y449, N450, L455, F456, T470, I472, E484, Y489, Q493, S494 |
| S107R | R346, K444, G447, Y449, N450, L455, E484, G485, F486, Y489, F490, L492, Q493, S494 |

## Supplementary Figures

**Supplementary Fig 1:**
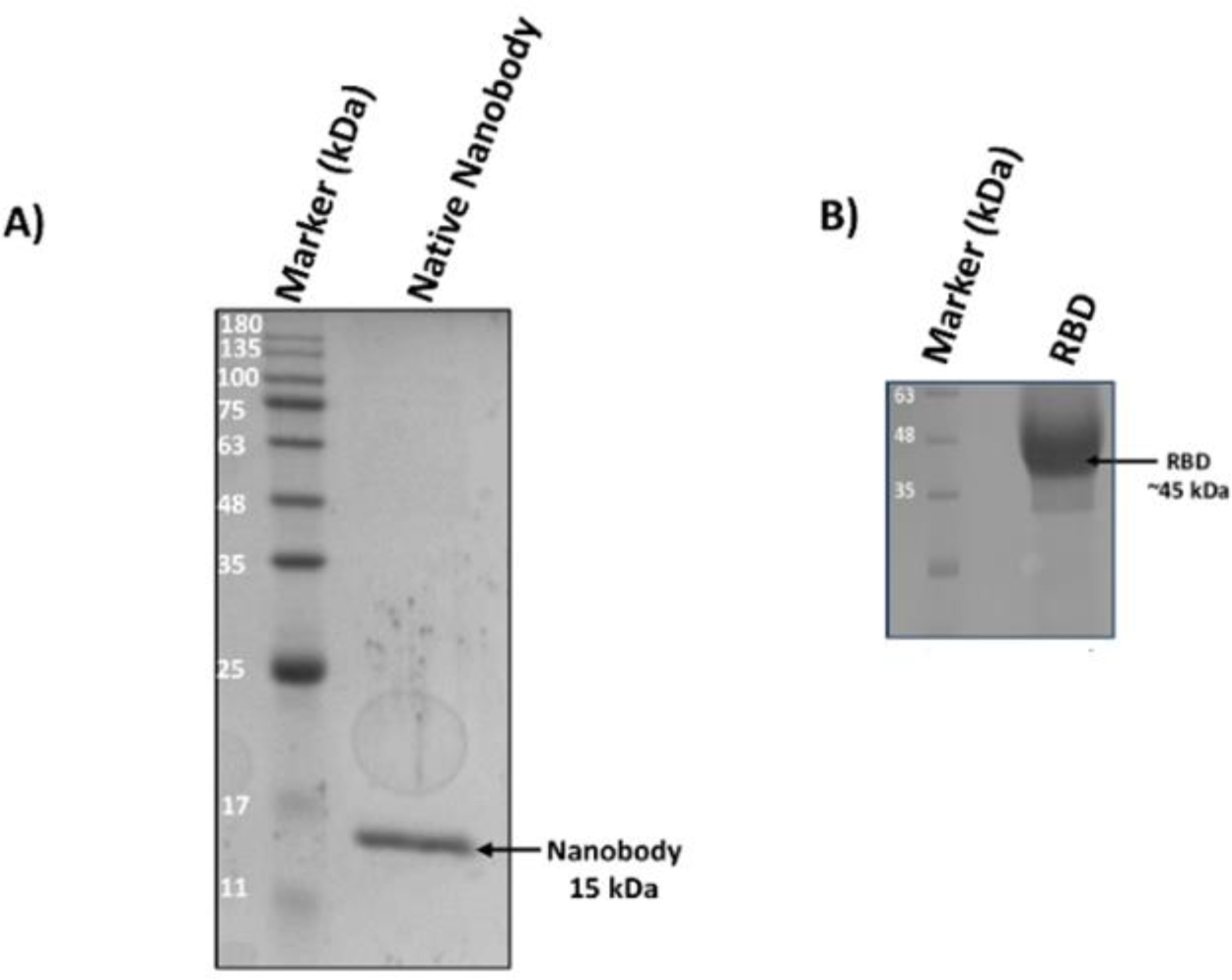
SDS-PAGE profile for the purification of native nanobody and RBD proteins. A) Lane 1: marker (180 kDa); Lane 2: native nanobody (15 kDa). B) Lane 1: marker; Lane 2: RBD (∼ 45 kDa).

**Supplementary Fig 2:**
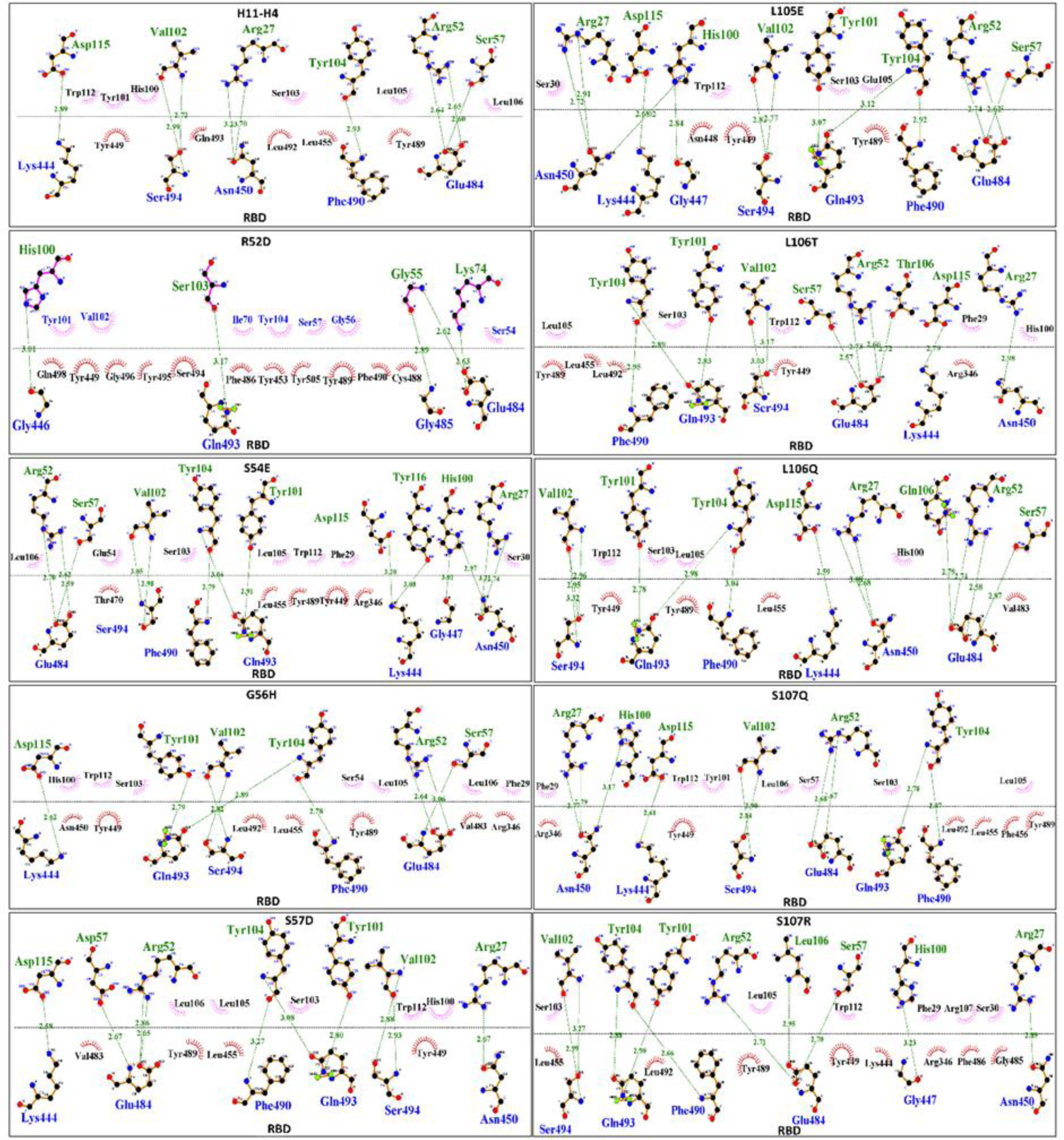
2D schematic representation of nanobody and RBD interacting residues generated through DimPlot. The docked complex of nanobody and RBD generated using HADDOCK 2.4 was used for the preparation of DimPlots. Hydrogen bonds are depicted by green dashed lines connecting the atoms involved, with the donor-acceptor distance indicated in Å. Hydrophobic contacts are shown as red arcs with spokes extending towards the atoms they interact with.

**Supplementary Fig 3:**
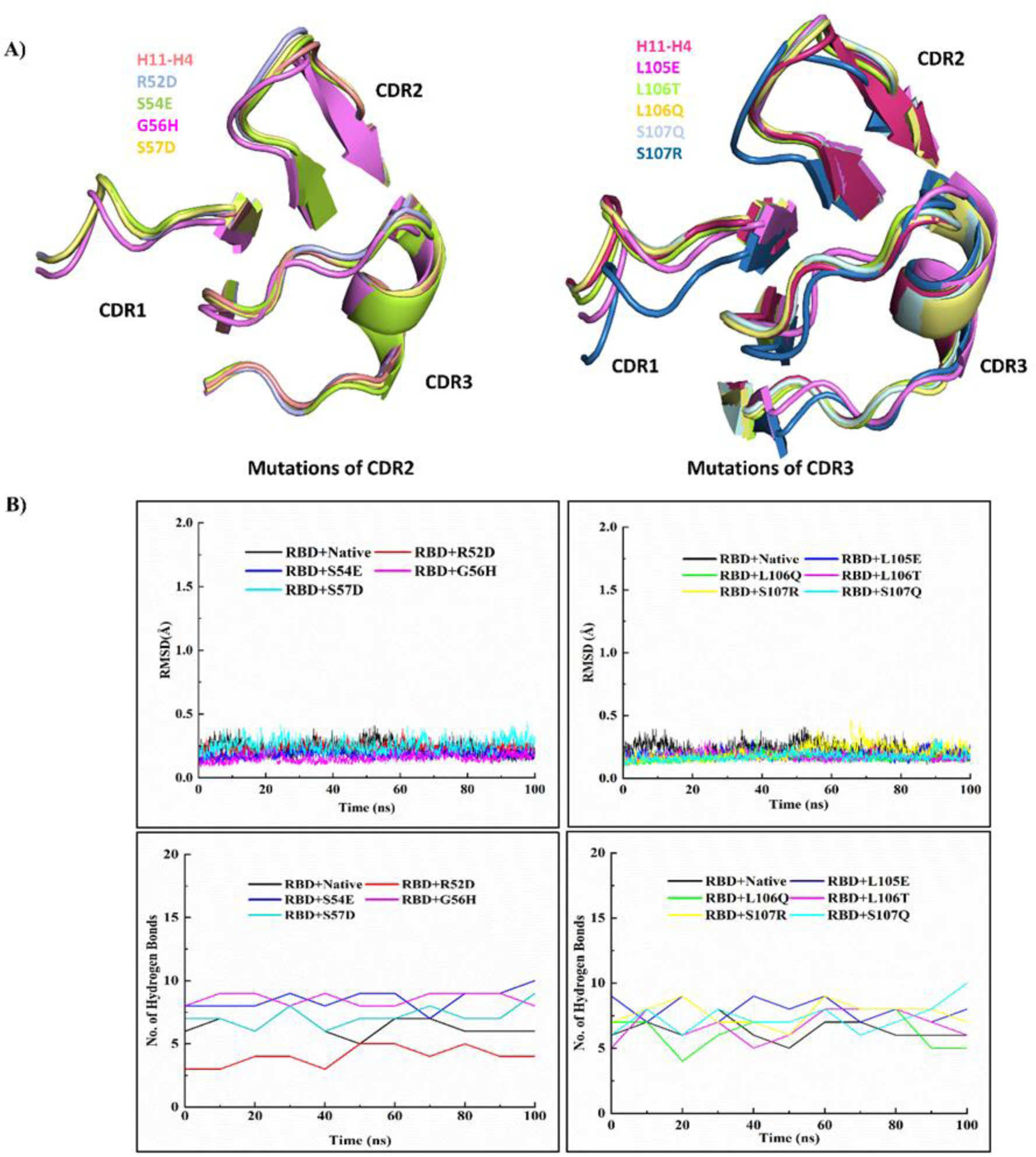
Structural superimposition of CDRs and analysis of MD simulations trajectories. A) CDRs of energy minimized structure of native and mutants for CDR2 and CDR3 were superimposed using PyMol. B) For the CDR2 and CDR3 the MD simulations trajectories were used to generate RMSD Plot for the native/mutant nanobody in complex with RBD and number of H-bonds formed between the native/mutant nanobody and RBD throughout 100 ns MD run.

**Supplementary Fig 4:**
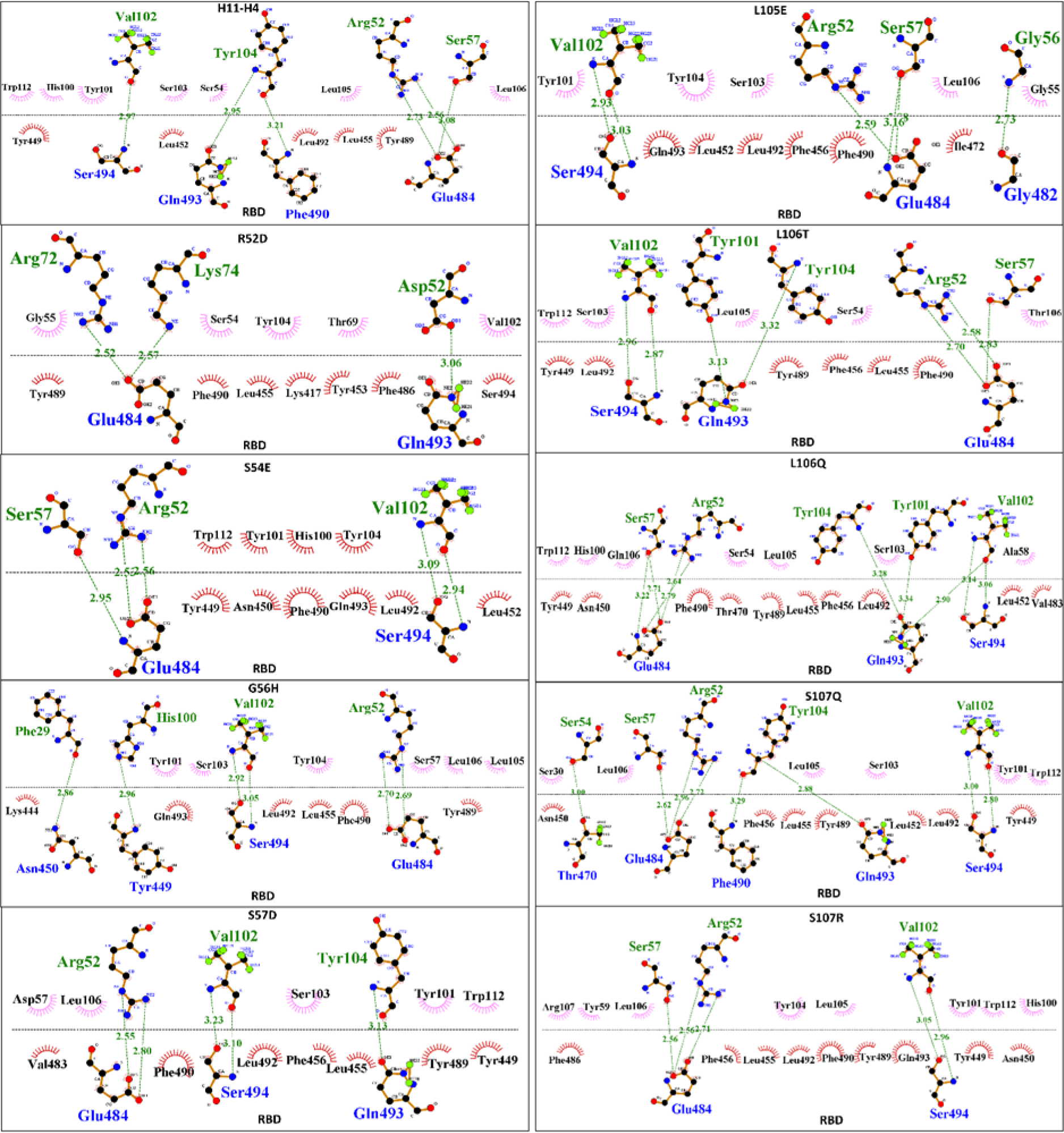
2D schematic representation of nanobody and RBD interacting residues generated through DimPlot. The frame PBDs were extracted from the MD run trajectories and used for the generation of DimPlots. Hydrogen bonds are depicted by green dashed lines connecting the atoms involved, with the donor-acceptor distance indicated in Å. Hydrophobic contacts are shown as red arcs with spokes extending towards the atoms they interact with.

